# Local and distributed information coding in the ventral stream

**DOI:** 10.1101/2025.03.09.642235

**Authors:** Paul C. Bogdan, Cortney Howard, Kirsten Gillette, Roberto Cabeza, Simon W. Davis

## Abstract

Neuroscience is replete with evidence that cognitive representations are distributed across many cortical regions. Yet, the scale and content of such distributed processing is unclear. Do findings of widespread information coding suggest a large-scale “forest” of regions interacting to represent information or instead imply a multitude of small-scale trees, processing information as localized modules. To investigate this distinction, we used visual and semantic representational analysis of fMRI data from 60 participants viewing everyday objects in multiple task contexts, and we examined the relationships between regions in terms of information coding. We demonstrate that coding of visual content in the occipital lobe is overwhelmingly modular, such that different occipital structures show limited coordination and tend to encode information redundantly. By contrast, the coding of semantic content in the inferior temporal lobe involves a high degree of coordination between regions, which optimize their coding to collectively represent a large semantic space with minimal redundancy between regions. No other brain area – neither the parietal nor prefrontal cortices – shows the preference for large-scale coding seen in the inferior temporal lobe. Taken together, these results outline a framework of how the ventral stream transitions from small-scale to large-scale coding as information progresses from visual to semantic representations.

**Significance statement:** How does the brain convert incoming signals into usable information? Many studies have investigated this question by attempting to clarify which brain regions encode what information (e.g., V4 encodes color information). We instead aimed to shed light on the degree of coordination among information coding regions. We find that the visual-to-semantic transition as information flows anteriorly in the occipitotemporal cortex is accompanied by a shift from modular to distributed coding. That is, occipital regions encode perceptual information relatively independently with redundancy, while inferior temporal lobe regions cooperate to most efficiently represent a large space of semantic information. By leveraging ideas from information theory, our work introduces coding scale as a new dimension for understanding the architecture of information coding.

## 1. Introduction

One of neuroscience’s oldest debates is whether the brain encodes information in a localized or distributed fashion. Early lesion studies were formative in localizing cognition to discrete brain regions,^1,2^ but even in early work, distributed processing schemes were sown with hypotheses on neurological disorders deriving from the disconnection of neural regions.^3–5^ Task-fMRI brought new tools to explore these ideas, and techniques like representational similarity analysis (RSA) have been used to identify dissociations between the regions encoding different types of information (e.g., visual or semantic).^67–9^ Furthermore, RSA has demonstrated how stimuli of all sorts may prompt information coding across a wide range of brain regions.^10–12^ However, it remains unclear whether such results on visual and semantic coding indicate a multitude of regions performing highly localized information processing or whether regions are interacting as part of some integrated system.

The ‘content’ of cortical representations can be summarized by two domains that dominate the cognitive neuroscience of object processing: *visual* features and *semantic* features, which seem to be most strongly represented in different brain areas. The cognitive neuroscience of vision has demonstrated this using convolutional neural networks (CNNs) to model visual processing along the occipitotemporal pathway. Early hidden layers reliably predict brain activity patterns predominantly in early visual cortex, while late layers predict activity in more anterior ventral pathways regions.^6,13^ Similarly, researchers focused on semantic cognition have shown how multiple loci in the anterior temporal cortex (perirhinal cortex, anterior temporal lobe) are associated with processing of multimodal object properties.^7–9^ These visual and semantic findings together imply a gradient of differential types of information processing, whereby more abstract semantic information is progressively encoded more anteriorly, but the scale of this well-known pattern are unclear.

The ‘scale’ in neural information processing also is quantifiable at multiple levels. In terms of abstract computation, a brain system may be highly modular with its subsets representing information in parallel and with fairly minimal interaction (**Figure 1A**). In terms of population coding, neural representations may be scattered across distant neurons interacting to encode information (**Figure 1B**). Questions concerning the scale of regional coordination in the representation of information can be investigated by (i) examining the amount of information expressed by a collection of regions together or by each region separately and (ii) measuring the redundancy in information expressed across regions.^14^ Compared to an integrated brain system that represents information with large-scale population codes, a more modular system that represents information in small specialized population codes has a lower capacity to encode information and will display greater redundancy between population codes. This link between specialization and redundancy may seem unintuitive, but it follows from the idea that a set of neurons can encode the maximum quantity of information if unconstrained and will avoid dedicating resources to duplicate coding (i.e., the Efficient Coding Hypothesis^15^). As a more concrete example, V4 is known to encode color while V5 encodes motion. However, these structures will presumably also express some overlapping information, which would not be duplicated in a hypothetical V4-V5 combined structure.

**Figure 1.**
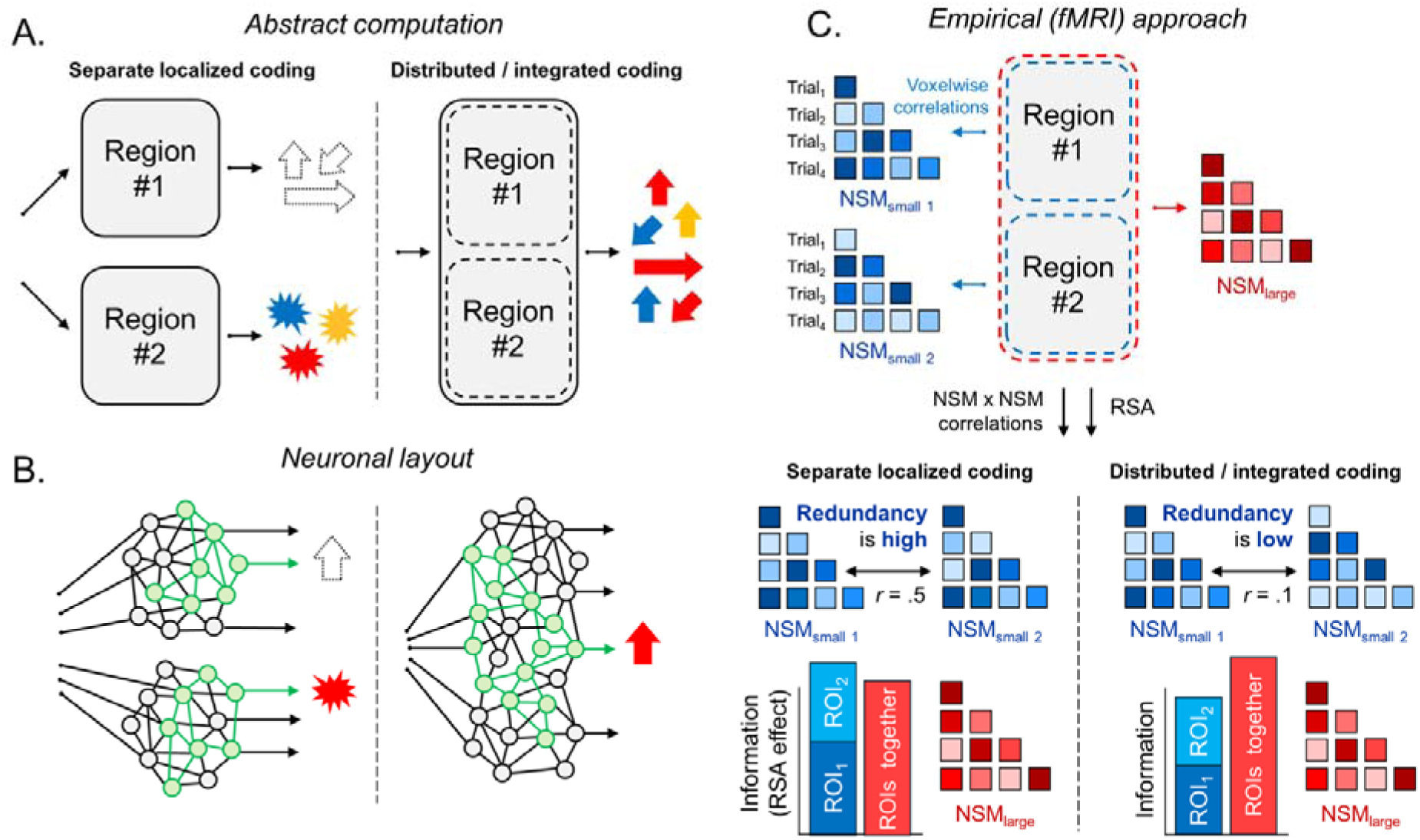
Illustration of the concepts targeted here. **A.** Different features can either be extracted separately or simultaneously, and regions can be conceptualized as encoding information as smaller modules or more integrated functional units. For example, Region 1 may code for motion, while Region 2 codes for color; alternatively, these regions may collectively encode both in an indivisible manner. **B.** Neuronally, features can be parsed separately via multiple small-scale population codes or simultaneously with larger-scale population codes. **C.** Representational similarity analysis (RSA) can be used to discern whether sets of regions encode information in a small- or large-scale manner. To model small-scale coding, neural similarity matrices (NSMs) can be extracted from trial-wise responses in regions separately (NSM_small 1_ & NSM_small 2_) or while pooling all voxels in a larger region (NSM_large_). We can test Information Processing by submitting these RSMs to RSA to evaluate the quantities of information encoded at each level. We can also test Redundancy by computing the correlations between small-scale NSMs; strong correlations between NSMs indicates redundant information coding. **Supplemental Materials 1** provides simulations describing how small/large-scale information coding may elicit different fMRI effects on decoding visual and semantic information, and demonstrates how RSA may distinguish these scales.

Scale in information coding can be empirically studied using fMRI and RSA, such that the correspondence between a theoretical representational similarity matrix (RSM) and a brain region’s neural similarity matrix (NSM) indicates that the information described in the RSM is represented by a brain area.^16,17^ To study whether regions coordinate synergistically to form a large-scale unit, the NSM can be defined while pooling voxels across regions (**Figure 1C**). If the resulting NSM better fits the theorized RSM better than expected from analyzing the regions individually, this suggests that the regions are coordinating their information processing (**Figure 1C**). Correlating just small-scale NSMs further allows measuring the extent pairs of regions encode similar information. Weak correlations would indicate minimal redundancy between regions, which can further point to regions operating as an integrated unit.

In sum, the organizing principles behind how the factors of scale and content are differentially coordinated across the brain are unknown. One possibility is that information coding in many regions reflects the coordination of a specific type of information to form an integrated computational whole that produces a large-scale population code. Alternatively, widespread information coding may reflect the coordination of multitudes of self-contained modules that operate mostly independently, differing in the specific type of information they encode but with possibly high redundancy. A large integrated system may be more efficient, but the inherent complexity may create challenges. Our research treats this type of information processing ‘scale’ as a possible property of brain systems (e.g., the occipital lobe, inferior temporal lobe, etc.), and examines how the scale of processing may depend on whether regions are encoding visual or semantic content.

Our research adapts traditional RSA techniques to study how collections of regions encode visual and semantic information in small- or large-scale fashions. Preliminary simulations formalize the claims above on “small-scale” and “large-scale” coding and demonstrate how these conditions can be distinguished using RSA (**Supplemental Materials 1**). Subsequently, our study uses fMRI data (N = 60 subjects) from a memory paradigm where 114 real-world objects were repeatedly shown across four stages involving (i) passive naming, (ii) contextual encoding, (iii) visual recognition, and (iv) conceptual recognition.^1^ The range of task demands across this dataset is explicitly incorporated to facilitate generalization of visual and semantic information across different task demands. Analyses were as follows: *First*, we examined the small versus large by investigating how pooling voxels from multiple regions may magnify RSA effects. This analysis is performed across different brain areas and for either perceptual or semantic information. *Second*, we measured correlations between regions to study redundancy in the information encoded in each region. *Third*, we assessed the relationship between these two phenomena, testing whether high collective information within a large-scale area is linked to redundancy among constituent regions.

Taken together, these analyses aim to provide support for the overarching perspective that understanding the scale of information processing it the key to understanding how different forms of information are decoded in the brain.

## 2. Results

### 2.1. Standard representational similarity analysis

Before distinguishing areas along the small/large distinction, we performed a traditional RSA analysis as a reference point. We parcellated the brain into 210 neocortical ROIs using the Brainnetome Atlas (mean size = 4.9 cm^3^). For each ROI, an NSM (Neural Similarity Matrix) was generated via across-trial correlations of the ROI’s voxels. Each ROI’s NSM was then correlated with a model RSM corresponding to perceptual features (early convolutional neural network activation) or semantic features (word2vec embedding). Next, second-order RSM *×* NSM_ROI_ correlations were computed separately for each participant and each of the experiment’s four stages. The correlation coefficients were averaged across stages by participant and submitted to one-sample t-tests for the group-level analysis (**Figure 2**). Perceptual information was overwhelmingly represented in posterior areas, while semantic representations were more anterior and spatially diffuse – consistent with earlier work.^8,18,19^ However, more elaborate analyses are needed to disentangle whether these patterns within individual ROIs wholly capture population codes or just components of larger ones.

**Figure 2.**
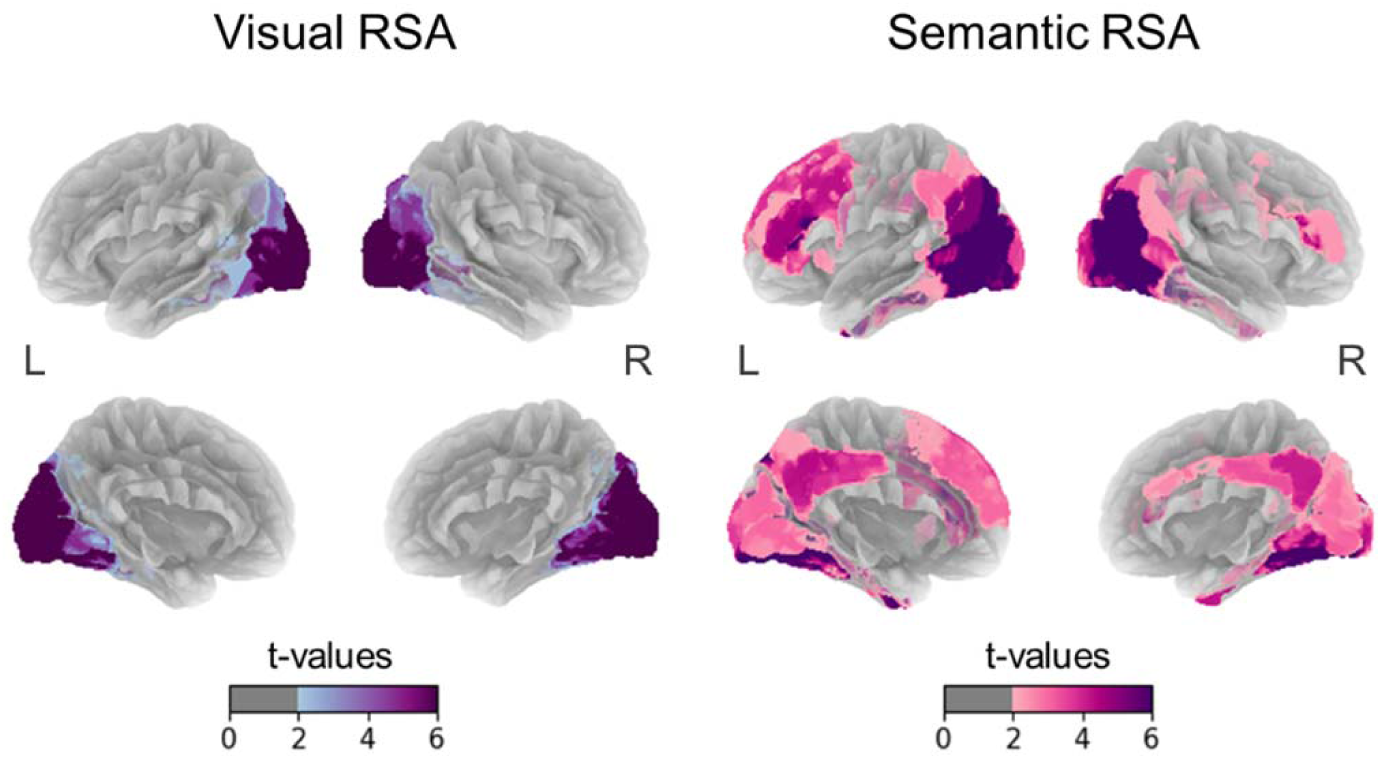
Traditional RSA results for perceptual or semantic coding. RSM (perceptual or semantic) × NSM_ROI_ correlations were performed separately for 210 neocortical. After averaging the correlation coefficients across the four stages, the data were submitted to one-sample t-tests whose results are shown here. Although 210 ROIs were tested, no correction for multiple comparisons in significance testing was applied, as these maps are simply meant as a reference for the later analyses.

### 2.2. Large and small-scale RSA

Our next analysis examined small-scale and large-scale coding across four areas: the occipital lobe, inferior temporal lobe, parietal lobe, and prefrontal cortex (PFC) (volume range = 138-288 cm^3^). We contrasted RSA effects between (i) a baseline NSM capturing the within-ROI coding, and (ii) an NSM that also accounts for large-scale codes spanning multiple ROIs. For an area’s baseline NSM, we averaged its constituent ROIs’ NSMs to form NSM_m-small_ (**Figure 3A**). Then, to model synergistic relationships, the voxels from its multiple ROIs were pooled (effectively producing one huge ROI), and voxel-wise correlations were used to define NSM_large_. The two NSMs were then separately submitted to second-order correlations. For the results, if an area’s NSM_large_ better tracks a model RSM, this would indicate that the area holistically encodes more information than the sum of its parts.

**Figure 3.**
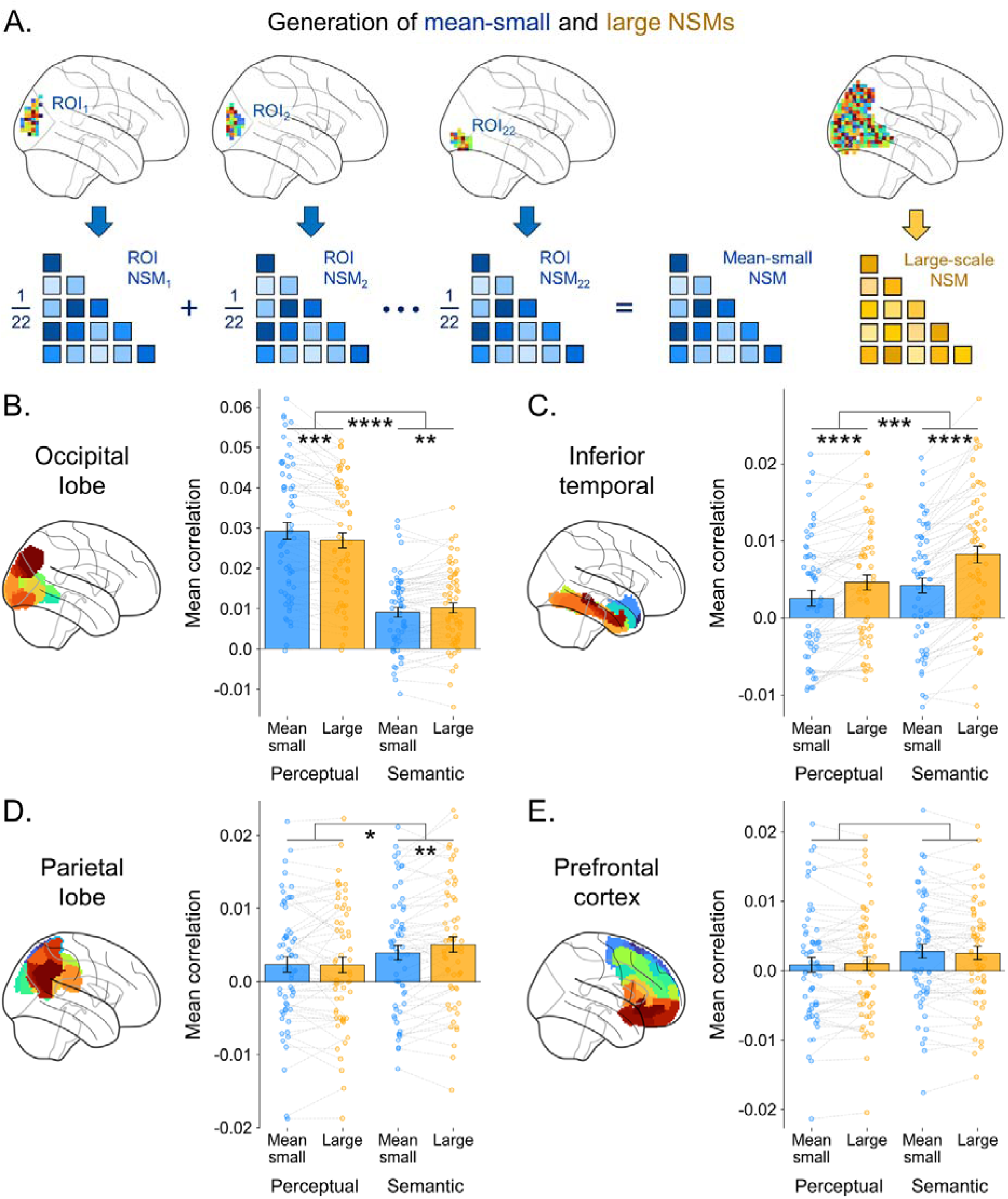
RSA results for small or large-scale coding. **A.** Illustration of the methodology used to generate the small/large-scale NSMs. This example is based on the 22 occipital ROIs. **B-E.** Results of the RSM (perceptual or semantic) × NSM (small or large) correlations for the four large areas tested. Dots represent individual participants, and the dashed lines indicate pairs from the same participant. Significance stars reflect paired t-tests or interactions from repeated-measure analyses of variance (ANOVAs). *, p < .05; **, p < .01; ***, p < .001; ****, p < .0001.

The occipital lobe emerged as a core for processing visual features but showed no signs of encoding information in a large-scale fashion (**Figure 3B**). In fact, particularly strong RSA effects emerged specifically for small-scale coding, suggesting the occipital lobe was characterized by particularly localized modules. This is a double dissociation relative to the inferior temporal lobe; the two-way Area (occipital/inferior temporal) × Scale (small/large) interaction for perceptual information is significant (*F*[1, 59] = 42.89, *p* < .0001). By contrast, the strongest inferior temporal lobe effects were seen for semantic representation and large-scale coding (**Figure 3C**). Measuring the Area × Scale interaction now for semantic information is again significant (*F*[1, 59] = 32.69, *p* < .0001). In the parietal lobe and prefrontal cortex, differences between coding sizes were smaller or non-existent (**Figures 3D & 3E**), pointing to the uniqueness of the occipital lobe’s small-scale coding and the inferior temporal lobe’s large-scale coding.

Alternative analytic approaches affirm these results. We tested an alternative approach to semantic modeling – using the last layer of the VGG16 – and reproduced the above interaction, showing that the inferior temporal lobe encodes specifically semantic information in a large-scale fashion (**Supplemental Materials 2**). Next, to potentially better disentangle small-scale from large-scale effects, we revised the large-scale NSM to be based on across-trial correlations between ROIs’ average activations (i.e., effectively treating each ROI as a single voxel; **Supplemental Materials 3**). This approach reproduced the prior interaction effects above on small-scale occipital representations of perceptual information and large-scale inferior temporal representations of semantic information. Additional analyses demonstrated significant large-scale coding effects in the inferior temporal lobe, even when regressing single ROI’s small-scale NSM separately, establishing that the wider patterns must reflect interactions between regions (**Supplemental Materials 4**).

Further analyses showed how large-scale semantic coding effects linked to the inferior temporal generally also emerge when examining the present study’s task stages individually rather than pooling their data (**Supplemental Materials 5**). Altogether, the findings consistently suggest that the inferior temporal lobe is organized such that relatively far-flung neuronal elements coordinate to represent semantic information as an integrated unit.

### 2.3. Redundancy in each area’s coding

The next stage of our analysis sought to address regional redundancy in visual and semantic processing. If a set of regions form integrated computational units with large-scale population codes, then it is expected that they will organize to minimize redundancy in their information processing, per the efficient coding hypothesis.^15^ To test this hypothesis, we addressed regional redundancy for representational, neural pattern, and similarity across time (i.e., “functional connectivity”). We first computed *NSM_ROI_ × NSM_ROI_* correlations between ROIs,^11^ which the degree that two regions differentiate the same pairs of objects. Each pair of ROIs’ correlations were computed for all four task stages then averaged. Compared to the mean representational similarity among occipital (mean *r* = .36 [.34, .38]), parietal (*r* = .33 [.32, .34]) and PFC (*r* = .31 [.30, .33]) regions, representational similarity in the inferior temporal lobe was markedly decreased (*r* = .26 [.25, .28]; Cohen’s *d* = −1.9 in comparison to the occipital lobe). These results suggest that inferior temporal regions display uniquely low levels of redundancy in information coding (**Figure 4A**).

**Figure 4.**
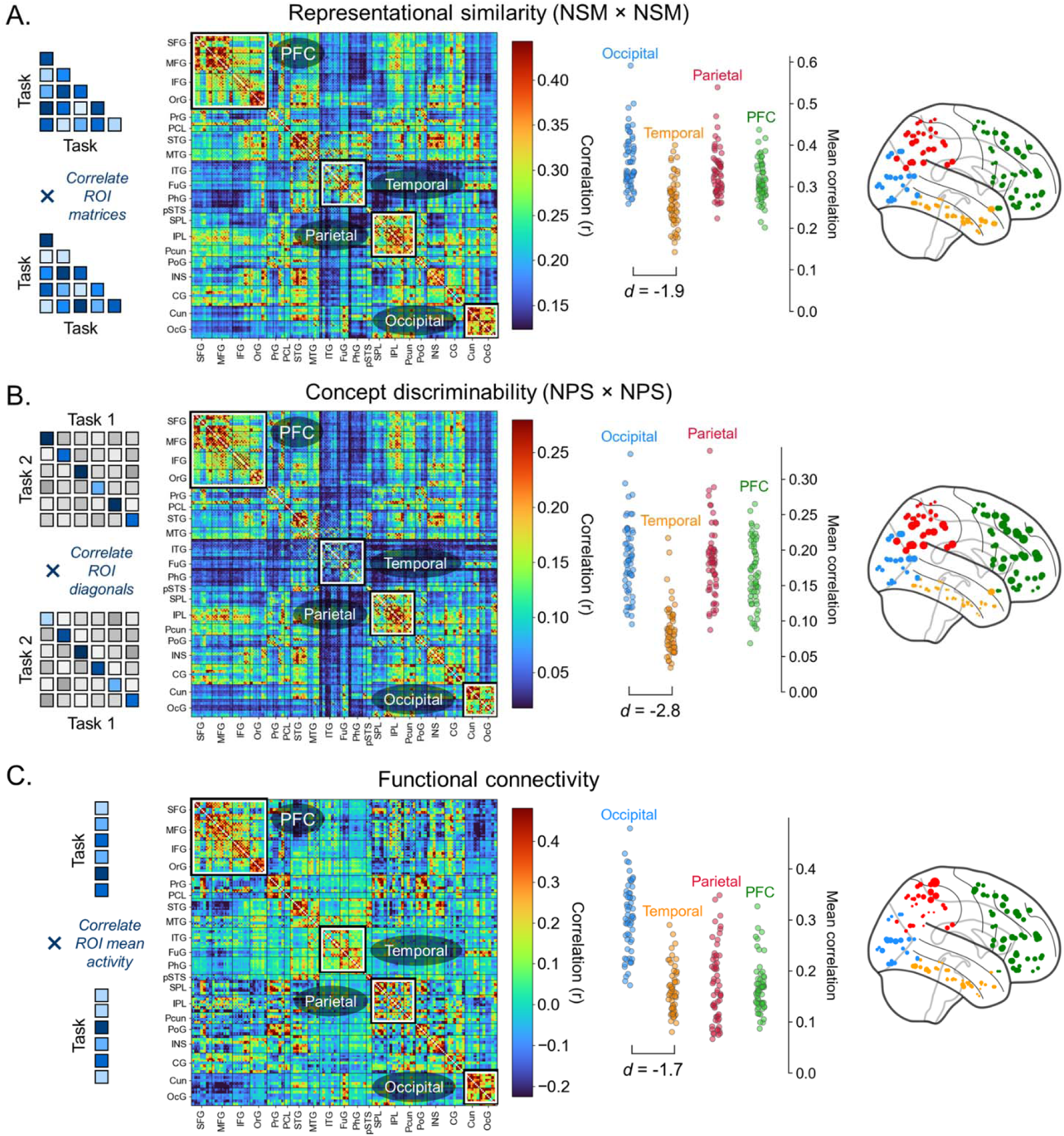
Heterogeneity in temporal lobe processing. The three rows represent ROI-ROI correlations between (**A.**) small-scale NSMs, correlated separately for the four tasks then averaged, (**B.**) neural pattern similarity (NPS) measures, computed and correlated separately for the six possible pairs of two tasks, then averaged (**C.**) functional connectivity, correlated separately for the four tasks then averaged; functional connectivity is not an information measure but is provided as a point of comparison Each dot in jitter plots next to each matrix represents one participant’s average correlation within the specified anatomical region. Cohen’s ds are reported from paired t-tests comparing the occipital and inferior temporal lobes. In the glass brains on the right, each marker corresponds to one ROI and the marker size represents the ROI’s mean correlation with its 10 physically closest neighbors in the same hemisphere.

Next, we exploit the multi-stage nature of our study to evaluate how different lobes may vary in the extent their constituent regions specialize in discriminating different concepts/concepts. This analysis adapts standard approaches to neural-pattern similarity analysis: If one ROI shows a consistent voxel-wise signature to a “cat” stimulus across a visual image of a cat and a verbal cue “cat”, then it can be said the ROI represents the concept “cat”. If said ROI does not show a consistent signature to a “house” stimulus but another ROI does, this indicates that the two ROIs are specialized for discriminating different concepts. We formally tested this by preparing an item-wise measure of neural-pattern similarity for each ROI (averaged across the six possible pairs of the four task stages) and then correlating this measure across ROIs. As in the analysis of representational similarity above, the correlations were again strikingly lower among inferior temporal regions (**Figure 4B**). Hence, there is a heightened degree of regional specialization for concepts in the inferior temporal lobe, further evidence of uniquely low redundancy in inferior-temporal information processing.

Lastly, to address coordination of regions independent of information coding we computed similarity across time, (i.e., “functional connectivity”) by examining the correlation between ROIs’ mean univariate activity timeseries. In our framework, functional connectivity quantifies the similarity in the univariate fluctuations between regions but is not a measure of the information processing performed by these regions. Accordingly, this analysis yielded less pronounced inferior-temporal drops (**Figure 4C**), indicating that these other trends specifically concern information coding. Taken together, the relatively lower correlations in representational and task-general similarity within the inferior temporal lobe are consistent with the area performing coordinated large-scale computations specific to information processing.

### 2.4. Large-scale representation and low redundancy

The final analyses assessed more direct relationships between information processing and redundancy. We specifically examine how visual or semantic representations in the occipital lobe or inferior temporal lobe may be linked to the level of redundancy in these areas. Based on the results thus far, we expect that the strong large-scale semantic representation in the inferior temporal lobe is linked to lower redundancy among inferior-temporal regions’ coding. For each participant and each of the four study stages, we calculated each area’s large-scale RSA effects (**Figure 3C**) and degree of redundancy among its regions (mean *NSM_ROI_ × NSM_ROI_*; **Figure 4A**). Submitting these measures to a multilevel linear regression reveals that, in the inferior temporal lobe, large-scale representation of semantics is tied to lower redundancy (β = −.17 [−.30, −.04], *p* = .01; **Figure 5**).

**Figure 5.**
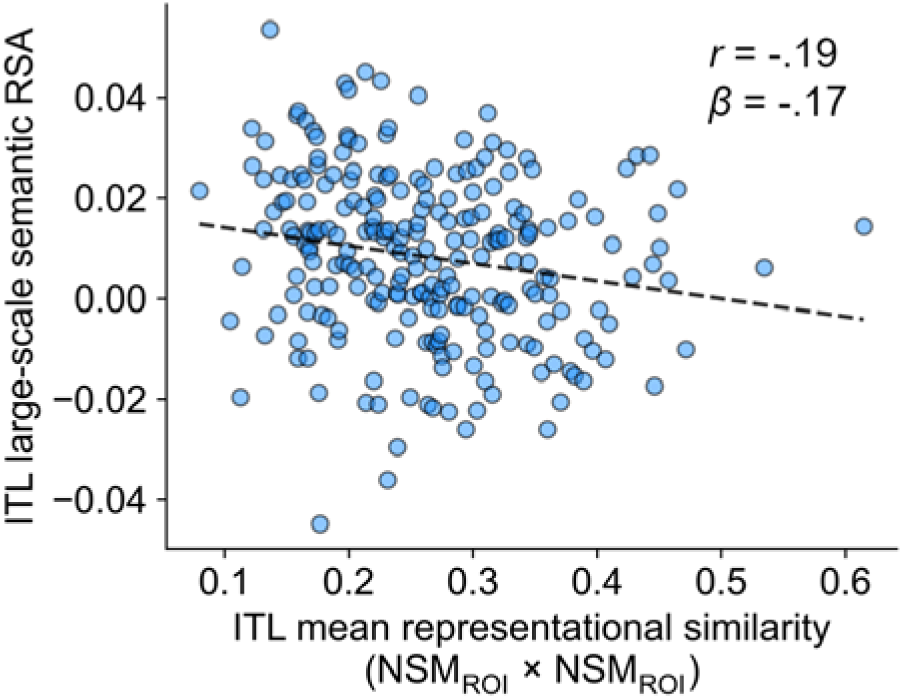
Assessing the link between distributed representation and regional correlations. Each dot corresponds to the data of one session from one participant. The x-axis corresponds to the mean NSM_ROI_ × NSM_ROI_ correlation, averaged across the inferior temporal lobe as in Figure 4A. The y-axis corresponds to the distributed RSA effect, computed as in Figure 3. The dashes show the negatively sloped line of best fit.

This effect is not significant when testing the cross-links between inferior-temporal redundancy and perceptual coding (β = −.06 [−.21, .09], *p* = .43), occipital redundancy and semantic coding (β = −.02 [−.06, .03], *p* = .50), or occipital redundancy and perceptual coding (β = .04 [.00, .08], *p* = .03; note, this effect is in the opposite direction). Hence, uniquely strong large-scale representation in the inferior temporal lobe relates to the area’s uniquely low level of redundancy between its regions’ information processing.

## 3. Discussion

The present research aimed to clarify the organization of computation and the size of population codes across the brain. The research yielded three conclusions. *First,* we performed RSA at different spatial scales, which indicated that, in the occipital lobe, perceptual processing is split across small-scale population codes. In contrast, inferior temporal information processing is distributed in a way not found anywhere else in the cortex, such that distant inferior temporal regions synergistically interact to encode semantic features. *Second*, we assessed the relationship between inferior temporal regions and found that the lobe is uniquely characterized by its constituent regions specializing in representing and distinguishing different objects. *Third*, we conducted analyses linking these different levels and showed that individual regions specializing in the information they encode promotes large-scale representation of semantic information. Overall, these results describe how the architecture of object processing in the ventral stream supports efficient information processing.

Perceptual information was most strongly represented in the occipital lobe and best modeled in terms of small-scale codes (**Figure 3**; supplemental materials **Figures S1-S3**). Our analyses defined perceptual information based on the first pooling layer of a convolutional neural network, which captures features like color, shape, and texture.^20^ Prior work has shown how these different aspects of visual function are relegated to distinct areas.^21,22^ Our results add that these distinct areas display relatively minimal coordination in information processing along with relatively high redundancy, pointing to a highly modular structure producing contained small-small population codes. This conclusion is consistent with lesioning research showing how V4 lesions destroy color processing but leave motion processing largely intact,^23^ while V5 lesions do the opposite.^24^ It is interesting to speculate on why small-scale coding may benefit the occipital lobe. In principle, more interconnected networks can encode more information.^25^ Yet, as the computer vision literature shows, fully connected layers are often sub-optimal at early stages, and restricted convolutional neurons ease training.^26^ In the human brain, a predisposition for modularity in early vision may similarly benefit perceptual development.

Contrasting the occipital patterns, our RSA results on semantic representation point to large-scale coding in the inferior temporal lobe (**Figures 3**; supplemental materials **Figures S1-S3**). That is, information coding is organized such that different inferior temporal lobes coordinate to efficiently represent semantic information. Interestingly, prior work mapping distributed coding of semantics has more often focused on the parietal, frontal, or lateral temporal lobes.^27–30^ Further, research on how information is linked across sensory modalities has found limited inferior temporal patterns.^31–33^ Yet, when our analyses found that the inferior temporal lobe displayed the strongest large-scale coding. As other regions largely did not show heightened large-scale coding effects, this may be a powerful angle through which to see the inferior temporal lobe specifically. Potentially, this functionality relates to the inferior temporal lobe’s central role in semantic memory encoding and retrieval. In general, a large set of neurons may better support a large semantic space than multiple smaller sets of neurons,^34^ and these efficiency gains may be particularly valuable for memory.

We additionally analyzed how a large-scale-coding system may organize its constituents to minimize redundancy. This involves correlating the representational signatures of neighboring regions (**Figure 4**; Supplemental Materials **Figure S4**). Within the inferior temporal lobe, NSM × NSM and NPS × NPS correlations are low between regions – markedly decreased compared to any other brain system. That is, inferior temporal ROIs highly vary in which objects they predominantly encode (NPS results) and which pairs of objects they predominantly distinguish (NSM results). This speaks to the synergy results on the inferior temporal lobe functioning as one or a small number of large computational units with great interconnection. Additionally, this finding aligns with prominent theories of inferior temporal organization focusing on its many category-specialized structures – e.g., for faces, locations, body parts, numbers, and letters.^35,36^ Our stimuli were diverse and not organized around known categories, and the present results suggest heterogeneity and specialization are general properties of the inferior temporal lobe.

These ROI-level properties support synergistic interactions and larger-scale representation per our final correlations between these levels (**Figure 5**). Our “integration account” of the inferior temporal lobe is not at odds with existing category-specialization theories. Instead, these are different sides of the same coin. For instance, rather than viewing the FFA and PPA as independently processing faces and locations respectively, we can conceptualize them as coordinating to encode both faces and places collectively. This is a special form of specialization, which differs from how the occipital lobe processes information in small population codes. Indeed, whereas the V4’s involvement in color coding and V5’s involvement in motion processing are narrow, the FFA plays a rich role in processing objects more generally.^37,38^ This is consistent with the FFA contributing to collective representations rather than operating in isolation.

In summary, the brain is a highly distributed representational system. Yet, the nature of this distribution varies according to tractable spatial and informational patterns. In some areas, information processing is segmented across regions that operate fairly independently. Other areas function more as synergistic collectives during information processing, where the loss of a region greatly hinders a large-scale population code. This dimension is particularly valuable for understanding the ventral stream. Incoming information begins as small perceptual codes in the occipital lobe and advances into large-scale semantic representations in the inferior temporal lobe. Future studies may also consider how representations may not be entirely organized into contiguous voxels but may emerge from synergistic interactions within a spatially disparate (non-contiguous) network. Further studies may also benefit mapping distribution more precisely than just along small or large scales, potentially developing atlases based on information synergy and redundancy properties. Broadly, the brain varies in the scale of its computational units, and understanding this dimension is key to seeing how neural structures promote efficient information coding.

## 4. Materials and methods

### 4.1. Participants

For this four-stage study, participants were recruited from the local community and screened to ensure they were native or fluent English speakers and no history of brain injury or mild cognitive impairment. In total, 38 younger adults (aged 18-30) and 38 older adults (aged 65-85) participated in the study. Of these, 16 people did not complete at least one of the four scans – e.g., due to withdrawing from the study after the first stage. The final set of 60 participants who provided fMRI data for all four stages consisted of a younger adult subset (N = 32, M_age_ = 22.8 [SD = 3.3], 66% female, 34% male) and an older adult subset (N = 28, M_age_ = 71.5 [SD = 4.5], 64% female, 36% male). There were no exclusions. The research was approved by the Duke University Institutional Review Board.

### 4.2. Task design

We rely on fMRI data collected during each task of a multi-stage experiment on how humans encode conceptual and relational information. The stages consisted of: (1) an object naming task, (2) a task where participants judged the link between an object and a scene, (3) a conceptual recognition task for object memory, and (4) a visual recognition task for object memory. Our analyses pool data across the four stages to draw conclusions that generalize across processing demands. In addition, pooling data was expected to increase the statistical power of our analyses, allowing robust conclusions. **Supplemental Materials 7.1** describes these four tasks in detail.

Additionally, **Supplemental Materials 2** shows the results of the small/large-scale RSA analysis using each task individually, showing patterns consistent with the main text conclusions.

### 4.3. Representational similarity analysis

MRI acquisition, preprocessing, and single-trial activation (beta) modeling procedures are detailed in **Supplemental Materials 7.2.** The procedures generated one beta value for each voxel and each trial, representing the intensity of the neural activation. These voxel-wise and trial-wise activations were used for RSA.

#### 4.3.1. Generating the model-based representational similarity matrices

RSA was conducted with respect to both visual and semantic features by defining a model RSM for each of these two types of information. The creation of each model RSM involved defining a visual or semantic feature vector for each of 114 total objects. To produce the visual vector for a given trial, every object image was submitted to the VGG16 convolutional neural network.^20^ For each object, all the activation values from neurons of the first VGG16 layer were extracted. For each object, its activation values (224×224×64 values) were flattened into a single vector (length = 3,211,264). These object vectors were reduced to smaller vectors of 114 values using PCA. Then, the visual RSM was generated via across-trial correlations of these vectors. For the semantic RSM, each trial’s vector was generated by submitting its object’s label to a pre-trained word2vec model;^39^ the public “word2vec-google-news-300” version from the *Gensim* Python package. The word2vec model yielded a 300-length vector for each of the same 114 objects, which was correlated between trials to generate the semantic RSM.

#### 4.3.2. Computing neural similarity matrices

All of the RSA performed in the report was ROI-based. Analyses parcellated the brain into 210 neocortical ROIs using the Brainnetome Atlas.^40^ The analyses also involved organizing a subset of the ROIs into occipital lobe, inferior temporal lobe, parietal lobe, and prefrontal cortex areas; see **Supplemental Materials 7.3** for information on each area’s constituent region names. The regional or area-wide voxel-wise data was used to generate NSMs via across-trial correlations. Second-order RSM × NSM Spearman correlations were computed for the different model RSMs and NSMs, done separately for each participant and each of the task stages. The correlation coefficients were then averaged by participant across task stages and then submitted to group-level analyses (paired t-tests or repeated-measures ANOVAs).

### 4.4. Redundancy analysis correlations

We pursued a series of analyses based on ROI-to-ROI relationships to examine patterns linked to when multiple ROIs encode overlapping/redundant information. We also sought to evaluate how this perspective compares to traditional functional connectivity analyses to confirm this approach adds new information above and beyond traditional means of assessing region-to-region interactions. First, analyses examined the similarity in small-scale NSMs, correlating these between ROIs – also referred to as model-free representational connectivity.^11^ The ROI-ROI correlations were calculated for each of the four stages and averaged. Second, analyses examined item specializing by calculating item-wise neural pattern similarity for each trial (also called the exemplar discriminability index);^41^ computing the neural pattern similarity of an ROI for a given object involved measuring the similarity of the voxel-wise response across two stages, then subtracting the average level of similarity (average correlation between the response to the object in one stage and the response to every other object in another stage). Computing neural pattern similarity for each object yielded a 114-length vector per ROI, as there were 114 objects in the task. This vector was correlated between ROIs. Third, analyses examined functional connectivity as the correlation between ROI’s trial-wise activations (averaged across voxels); often referred to as beta series functional connectivity.^42^

### 4.5. Linking large-scale representation and ROI-ROI correlations

The final set of analyses sought to link those representational scale to ROI-level information processing. For this, the analyses used the *NSM_ROI_ × NSM_ROI_* correlation matrices generated just above. For each participant and each of the four stages, the mean correlation was computed among ROIs within the occipital and inferior temporal lobes. For example, as there are 32 inferior temporal ROIs and thus 496 possible pairs of ROIs, the mean *NSM_ROI_ × NSM_ROI_* correlation amounts to averaging across those 496 pairs. Along with computing this mean, for each participant and each of the four stages, the degree of large-scale visual and semantic representation was also calculated.

This was done using the initial technique, which measured voxel-wise correlations while pooling every inferior temporal lobe ROIs’ voxels and then performed a second-order correlation with a semantic model RSM. Thus, for the 60 participants and four stages, one measure of the *NSM_ROI_ × NSM_ROI_* correlations and one measure of the degree of large-scale representation were computed. These 240 data points were submitted to a multilevel regression, linking the two measures: *Large scale RSA ∼ 1 + ROI corr + (1 | participant)*. For the analysis of the occipital lobe, *ROI corr* was strongly linked to “*small scale RSA*” for visual and semantic coding (*p*s < .001), and the degree of small scale RSA effects was added as a covariate. The significance of the fixed effect linking the measures was evaluated using the *lmerTest* package.^43^

## Data availability

The data collected will be shared upon request via email to the corresponding author, Paul C. Bogdan, or principal investigators Roberto Cabeza and Simon W. Davis; due to IRB restrictions, the data cannot be uploaded to a public repository. Code used to compile the results will be uploaded to a public repository prior to publication.

## Footnotes

1 In principle, our analyses could be done with any dataset where participants viewed diverse stimuli, but we opted for this data already collected by our group, as our experiment included multiple stages, this increases our ability to generalize across processing demands. In addition, our dataset is large: roughly 120 hours of task-fMRI scanning. We expected that this large size would allow fine-grain comparisons between RSA effects with high statistical power.

## Notes

### Competing Interest Statement

The authors have declared no competing interest.

### Summary of Updates

Edits to the phrasing in parts of the manuscript

### Funding

- National Institute on Aging: R01-618AG066901, R01-AG075417

## References

1. Alexander, M. P., Naeser, M. A. & Palumbo, C. Broca’s area aphasias: aphasia after lesions including the frontal operculum. Neurology 40, 353–353 (1990).

2. Wernicke, C. The aphasic symptom-complex: a psychological study on an anatomical basis. Archives of Neurology 22, 280–282 (1970).

3. Catani, M. & Ffytche, D. H. The rises and falls of disconnection syndromes. Brain 128, 2224– 2239 (2005).

4. Geschwind, N. Disconnexion syndromes in animals and man. Brain 88, 585–585 (1965).

5. Mah, Y.-H., Husain, M., Rees, G. & Nachev, P. Human brain lesion-deficit inference remapped. Brain 137, 2522–2531 (2014).

6. Khaligh-Razavi, S.-M. & Kriegeskorte, N. Deep supervised, but not unsupervised, models may explain IT cortical representation. PLoS computational biology 10, e1003915 (2014).

7. Binney, R. J., Hoffman, P. & Lambon Ralph, M. A. Mapping the multiple graded contributions of the anterior temporal lobe representational hub to abstract and social concepts: evidence from distortion-corrected fMRI. Cerebral cortex 26, 4227–4241 (2016).

8. Clarke, A. & Tyler, L. K. Object-specific semantic coding in human perirhinal cortex. Journal of Neuroscience 34, 4766–4775 (2014).

9. Tyler, L. K. et al. Objects and categories: feature statistics and object processing in the ventral stream. Journal of cognitive neuroscience 25, 1723–1735 (2013).

10. Haxby, J. V., Connolly, A. C. & Guntupalli, J. S. Decoding neural representational spaces using multivariate pattern analysis. Annual review of neuroscience 37, 435–456 (2014).

11. Huang, S., De Brigard, F., Cabeza, R. & Davis, S. W. Connectivity analyses for task-based fMRI. Physics of Life Reviews (2024).

12. Williams, N. & Henson, R. N. Recent advances in functional neuroimaging analysis for cognitive neuroscience. Brain and Neuroscience Advances 2, 2398212817752727 (2018).

13. Wen, H. et al. Neural encoding and decoding with deep learning for dynamic natural vision. Cerebral cortex 28, 4136–4160 (2018).

14. Luppi, A. I., Rosas, F. E., Mediano, P. A., Menon, D. K. & Stamatakis, E. A. Information decomposition and the informational architecture of the brain. Trends in Cognitive Sciences (2024).

15. Barlow, H. B. Possible principles underlying the transformation of sensory messages. Sensory communication 1, 217–233 (1961).

16. Kriegeskorte, N., Mur, M. & Bandettini, P. A. Representational similarity analysis-connecting the branches of systems neuroscience. Frontiers in systems neuroscience 2, 4 (2008).

17. Kriegeskorte, N., Cusack, R. & Bandettini, P. How does an fMRI voxel sample the neuronal activity pattern: compact-kernel or complex spatiotemporal filter? Neuroimage 49, 1965–1976 (2010).

18. Davis, S. W. et al. Visual and semantic representations predict subsequent memory in perceptual and conceptual memory tests. Cerebral Cortex 31, 974–992 (2021).

19. Martin, C. B., Douglas, D., Newsome, R. N., Man, L. L. & Barense, M. D. Integrative and distinctive coding of visual and conceptual object features in the ventral visual stream. elife 7, e31873 (2018).

20. Simonyan, K. Very deep convolutional networks for large-scale image recognition. arXiv preprint arXiv:1409.1556 (2014).

21. Cavina-Pratesi, C., Kentridge, R., Heywood, C. & Milner, A. Separate channels for processing form, texture, and color: evidence from fMRI adaptation and visual object agnosia. Cerebral cortex 20, 2319–2332 (2010).

22. Rolls, E. T., Deco, G., Huang, C.-C. & Feng, J. Multiple cortical visual streams in humans. Cerebral Cortex 33, 3319–3349 (2023).

23. Zeki, S. A century of cerebral achromatopsia. Brain 113, 1721–1777 (1990).

24. Newsome, W. T. & Pare, E. B. A selective impairment of motion perception following lesions of the middle temporal visual area (MT). Journal of Neuroscience 8, 2201–2211 (1988).

25. Roudi, Y. & Latham, P. E. A balanced memory network. PLoS computational biology 3, e141 (2007).

26. Krizhevsky, A., Sutskever, I. & Hinton, G. E. Imagenet classification with deep convolutional neural networks. Advances in neural information processing systems 25, (2012).

27. Carota, F., Nili, H., Pulvermüller, F. & Kriegeskorte, N. Distinct fronto-temporal substrates of distributional and taxonomic similarity among words: evidence from RSA of BOLD signals. NeuroImage 224, 117408 (2021).

28. Huth, A. G., De Heer, W. A., Griffiths, T. L., Theunissen, F. E. & Gallant, J. L. Natural speech reveals the semantic maps that tile human cerebral cortex. Nature 532, 453–458 (2016).

29. Kidder, A., Silson, E. H., Nau, M. & Baker, C. I. Distributed Cortical Regions for the Recall of People, Places, and Objects. eneuro 12, (2025).

30. Meersmans, K. et al. Representation of associative and affective semantic similarity of abstract words in the lateral temporal perisylvian language regions. Neuroimage 217, 116892 (2020).

31. Fernandino, L., Tong, J.-Q., Conant, L. L., Humphries, C. J. & Binder, J. R. Decoding the information structure underlying the neural representation of concepts. Proceedings of the National Academy of Sciences 119, e2108091119 (2022).

32. Fernandino, L. et al. Concept representation reflects multimodal abstraction: A framework for embodied semantics. Cerebral cortex 26, 2018–2034 (2016).

33. Tong, J. et al. A distributed network for multimodal experiential representation of concepts. Journal of Neuroscience 42, 7121–7130 (2022).

34. Elhage, N. et al. Toy models of superposition. arXiv preprint arXiv:2209.10652 (2022).

35. Kanwisher, N. Functional specificity in the human brain: a window into the functional architecture of the mind. Proceedings of the national academy of sciences 107, 11163–11170 (2010).

36. Pitcher, D., Charles, L., Devlin, J. T., Walsh, V. & Duchaine, B. Triple dissociation of faces, bodies, and objects in extrastriate cortex. Current Biology 19, 319–324 (2009).

37. Haist, F., Lee, K. & Stiles, J. Individuating faces and common objects produces equal responses in putative face-processing areas in the ventral occipitotemporal cortex. Frontiers in human neuroscience 4, 181 (2010).

38. Zachariou, V., Safiullah, Z. N. & Ungerleider, L. G. The fusiform and occipital face areas can process a nonface category equivalently to faces. Journal of cognitive neuroscience 30, 1499– 1516 (2018).

39. Mikolov, T., Chen, K., Corrado, G. & Dean, J. Efficient estimation of word representations in vector space. arXiv preprint arXiv:1301.3781 (2013).

40. Fan, L. et al. The human brainnetome atlas: a new brain atlas based on connectional architecture. Cerebral cortex 26, 3508–3526 (2016).

41. Nili, H., Walther, A., Alink, A. & Kriegeskorte, N. Inferring exemplar discriminability in brain representations. Plos one 15, e0232551 (2020).

42. Rissman, J., Gazzaley, A. & D’Esposito, M. Measuring functional connectivity during distinct stages of a cognitive task. Neuroimage 23, 752–763 (2004).

43. Kuznetsova, A., Brockhoff, P. B. & Christensen, R. H. B. lmerTest package: tests in linear mixed effects models. J Stat Softw 82, (2017).

