## Supplemental Materials for "Local and distributed information coding in the ventral stream"

### 1. Simulations of small and large-scale representation

We conducted simulations that formalize the concept of "large-scale" coding – stimulus representations being distributed across multiple regions coordinating to fully represent a stimulus space. In addition, we aim to demonstrate how the Representational Similarity Analysis (RSA) techniques that would be employed in the fMRI analyses serve to distinguish large-scale and small-scale coding scenarios.

The simulations suppose that items are composed of features, and each feature when active elicits some set of voxels in some region(s) to increase in activity. This setup permits the construction of a model RSM (based on the underlying feature data) and NSMs (based on the voxel data). A small-scale coding condition effectively serves as a baseline, where we assume that all regions all encode the same features and all regions encode information for all items (i.e., each region operates independently, and there is no coordination between regions). Large-scale coding is defined as a condition where different regions respond to different features or encode all features but only for different items. A mall-scale coding, by contrast, is treated as the baseline.

#### 1.1. Simulating features and brain data

More formally, it was supposed that there existed a set of possible binary semantic features that an item could have, designated as F (e.g., 50) possible features. There were I (e.g., 100) items, and each item was characterized by $N_F$ (e.g., 5) random features, represented as F-length one-hot vectors, each with $N_F$ ones and the remaining features as zeros. R (e.g., 50) regions are simulated, each comprised of V (e.g., 500) voxels. If a brain region was sensitive to a given feature, a certain proportion, P (e.g., 20%), of its voxels would activate with a value of +1. Random noise was also added to the activation of all voxels within each brain region (e.g., $\sigma = 25$ added to each voxel independently); the conclusions below do not hinge on the amount of noise added, we have simply picked this value and others below to generally produce the second-order correlations seen in brain data (e.g., $.01 < r < .05$).

**1.2. Representational similarity analysis**

Based on the simulated feature one-hot vectors, a model RSM was developed based on Pearson correlations between each item's one-hot vector. NSMs were computed to capture the overall degree small-scale or large-scale coding among all regions using a procedure that would be employed for the actual fMRI analysis. To define the NSM for overall small-scale coding, one $NSM_{region}$ was computed for each simulated region. All regions' $NSM_{region}$s were then averaged to create $NSM_{m\text{-}small}$. To define the NSM for large-scale coding, all of the regions' voxel-wise data were pooled, effectively forming a single large region. $NSM_{large}$ was then computed as based on across-trial correlations with respect to these pooled voxels.

**1.3. Experiments and results**

Several simulation designs were implemented to explore different scenarios of information coding. For each design, RSA was performed twice, measuring the Spearman correlation between the model RSM and either $NSM_{m\text{-}small}$ or $NSM_{large}$.

***Small-scale coding scenario:*** This experiment employed the parameters described in Section 1.1 above, simulating 50 regions that are each sensitive to every simulated feature. The second-order correlations between the model RSM and $NSM_{m\text{-}small}$ or $NSM_{large}$ were measured, then averaged across ten simulations. This yields similar second-order correlations for $NSM_{m\text{-}small}$ (mean $r$ = .047) and $NSM_{large}$ (mean $r$ = .047).

Here, we demonstrate just that small-scale coding leads to equal RSA effects across the two RSMs, although the fMRI results of the main text would ultimately demonstrate that in the occipital lobe, $NSM_{m\text{-}sall}$ surpasses $NSM_{large}$ in its RSA effect for perceptual information. One reason why this may have occurred is due to differences in the amount of noise between regions. As the preliminary single-ROI analyses show in the first Results subsection (Figure 2), some occipital ROIs tend to elicit stronger RSA effects than other occipital ROIs. In the small-scale-coding simulations, adding variable levels of noise to each ROI – adding σ = 25 noise to half of ROIs and σ = 100 noise to the other half – leads to $NSM_{m\text{-}sall}$ (mean $r$ = .014) surpassing $NSM_{large}$ (mean $r$ = .002) in RSA effects.

***Large-scale coding with feature distribution*:** This experiment modifies the small-scale coding scenario such that each region encodes just one feature; the fifty regions altogether encode the fifty features as so; the noise was also decreased to σ = 3 to accommodate for the loss in information compared to the small-scale coding experiment where every region was encoding all features. Here, the second-order correlations are higher for RSA using $NSM_{large}$ (mean $r$ = .047) than $NSM_{m\text{-}small}$ (mean $r$ = .038). Thus, contrasting RSA effects across these techniques successfully identifies this condition where regions are characterized by large-scale coding.

***Large-scale coding with item distribution*:** In this final simulation, each region was made sensitive to a unique, non-overlapping subset of items, such that the fifty regions were each sensitive to two items, collectively capturing all one hundred items; the noise was decreased further to σ = 0.1. Under this condition, second-order correlations are again higher for RSA using $NSM_{large}$ (mean $r$ = .007) than $NSM_{m\text{-}small}$ (mean $r$ = .000). Thus, contrasting RSA effects across these techniques successfully also identifies this condition where regions are characterized by large-scale coding.

We present here two possible manners by which large-scale information coding can be distributed across different regions: feature specialization or item specialization. Our RSA analyses do not attempt to distinguish between these possibilities, although in the fMRI analyses, the examination of redundancy (Section 2.3) will demonstrate that both large-scale coding formats are likely at play.

Altogether, these simulations formalize large-scale coding and show how contrasting NSMs in RSA can distinguish between whether brain regions are characterized by small- or large-scale coding. Our fMRI analyses (Section 2.2) would use the same analytical approach.

**2. Examining semantic representation using the last layer of VGG16**

Modeling semantic information using the last pooling layer of VGG16 (rather than word2vec, as used for every other analysis) yields large-scale coding RSA effects consistent with those found elsewhere in the present research. The occipital lobe continues to demonstrate a preference for encoding perceptual information and particularly through small-scale coding (**Figure S1A**), and the inferior temporal lobe continues to demonstrate a preference for encoding semantic information and particularly through large-scale coding (**Figure S1B**).

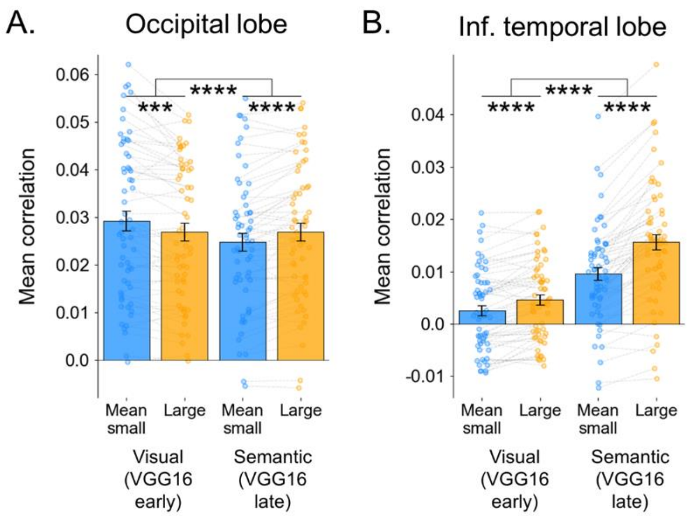


***Figure S1A. RSA results for small or large-scale coding using VGG16 to model semantics.*** *This figure uses the same analytic approach as for Figure 3 in the main text, but instead of using word2vec features to model semantic information, the final pooling layer of VGG16 is used. Notably, perceptual modeling is based on the first pooling layer of VGG16. ***, $p < .001$; ****, $p < .0001$.*

**3. Modeling large-scale coding via averaging**

Several further tests were done to confirm the large-scale coding effects in the inferior temporal lobe. The first test focused on disentangling RSA effects common to the within-ROI and area-wide results. The area-wide NSM was revised to be based on across-trial correlations between ROIs' average activations (i.e., essentially treating each ROI as one voxel). For example, there were 32 inferior temporal lobe ROIs, so the $NSM_{large\text{-}avgs}$ correlations were taken between 32-length vectors. Averaging by ROI eliminated any within-ROI patterns. Along with this change, the original RSM × NSM strategy was revised to regress the model RSMs on small-scale and large-scale NSMs together

($RSM \sim NSM_{m\text{-}small} + NSM_{large\text{-}avgs}$) (**Figure S2A**). This produced the same double dissociations as above with significant Area × Size interactions for perceptual information ($F[1, 59] = 32.15$, $p < .0001$) and significant Area × Size interactions for semantic information ($F[1, 59] = 10.31$, $p = .002$) (**Figures S2B & S2C**). There were no significant differences in the parietal lobe or PFC (**Figures S2D & S2E**), meaning that these changes in coding size specifically concern how the ventral stream.

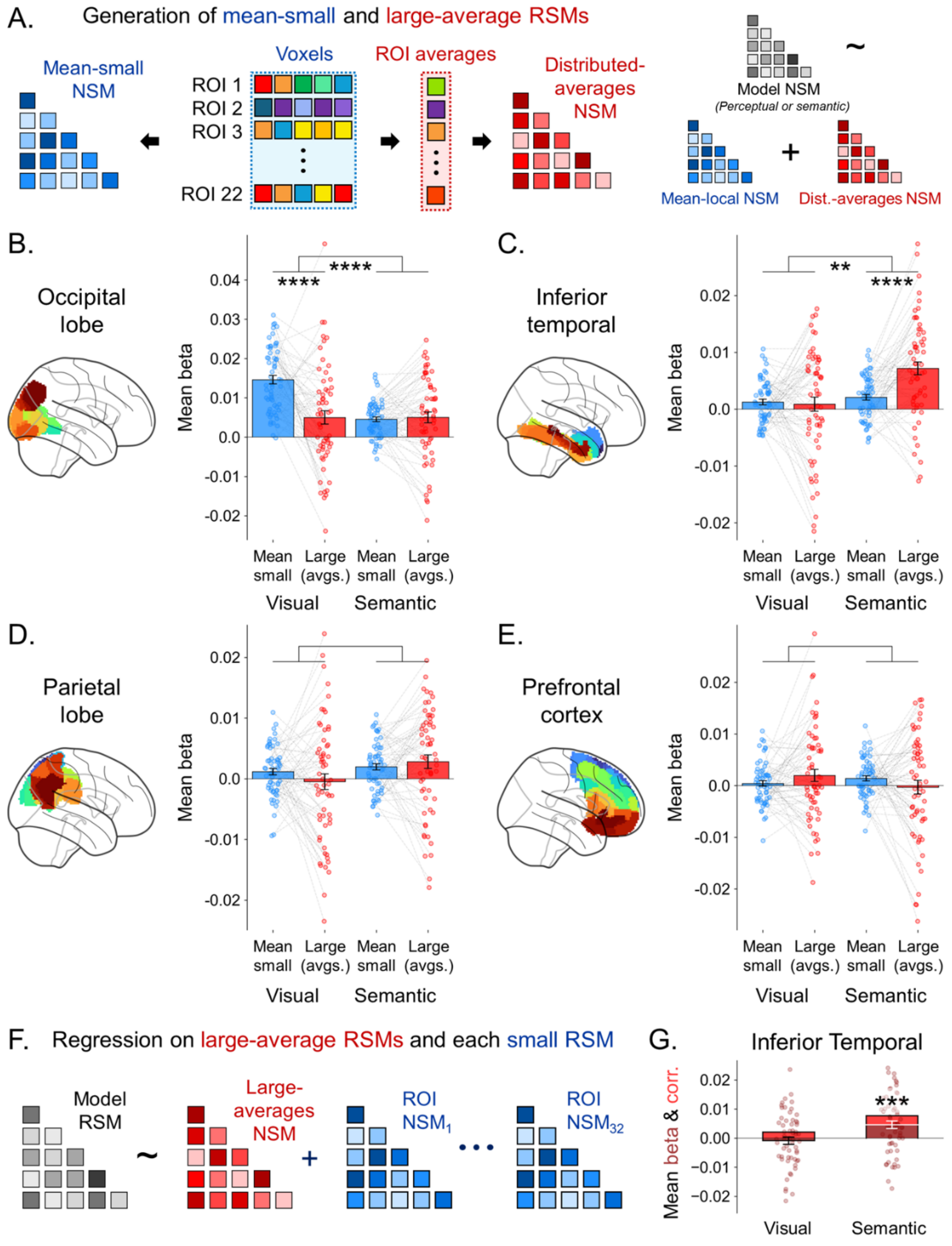

**Figure S2. RSA results for small-scale or large-scale (ROI averages) coding.** ***A.*** *Unlike the main text Figure 3 analyses, here a distributed NSM is defined with correlations between constituent ROIs' average activities. Also, whereas the Figures 2 and 3 methodologies performed RSA with RSM × NSM correlations, here it is done with a regression of RSM ~* $NSM_{m\text{-}small}$ + $NSM_{large\text{-}avgs}$. ***B-E****. Group-mean bars, participant dots, and paired t-test/ANOVA interaction results are reported as in Figure 3.* ***F.*** *A separate analysis focused on just the inferior temporal lobe and regressed a model RSM on the large-averages NSM along with all 32 inferior temporal lobe small-scale NSMs.* ***G.*** *The results of this many-predictor regression are shown in dark red. The three white stars indicate that the* $NSM_{large\text{-}avgs}$ *beta significantly surpasses zero. For reference, the results of RSM ×* $NSM_{large\text{-}avgs}$ *correlations are also shown in lighter red. *, p < .05; **, p < .01; ***, p < .001.*

### 4. Stricter multiple regression analysis

Yet stricter analyses specifically attempted to identify a distributed synergy effect in the inferior-temporal lobe while fully accounting for within-ROI coding. Here, model RSMs were regressed on $NSM_{large\text{-}avgs}$ along with all 32 constituent ROIs' small-scale NSMs ($RSM \sim NSM_{large\text{-}avgs} + NSM_{ROI\,1} + \ldots + NSM_{ROI\,32}$) (**Figure S2F**). Our focus is on the $NSM_{large\text{-}avgs}$ beta coefficient, and whether it surpasses zero while the other predictors ruled out small-scale coding in any one ROI. Group-level analyses indeed showed that the inferior temporal lobe's $NSM_{large\text{-}avgs}$ still significantly tracked semantic information (**Figure S2G**). This is strong evidence of large-scale coding – i.e., information encoded collectively that cannot be explained by the activity within any individual ROI.

### 5. Single-stage results

We performed the main text small/large-scale RSA comparison along with the above ROI-averaging based approach (for **Figure S2**), now examining each of the four task stages individually. The results are shown in **Figure S3** and are broadly consistent with main-text **Figure 3** and immediately above **Figure S2**. Note that there are some differences – e.g., the interaction effects are not significant in every case, or large-scale coding is also found to characterize inferior temporal lobe perceptual coding. However, these differences compared to the broader trends toward similarity are expected to be noise and not invalidate our primary conclusions.

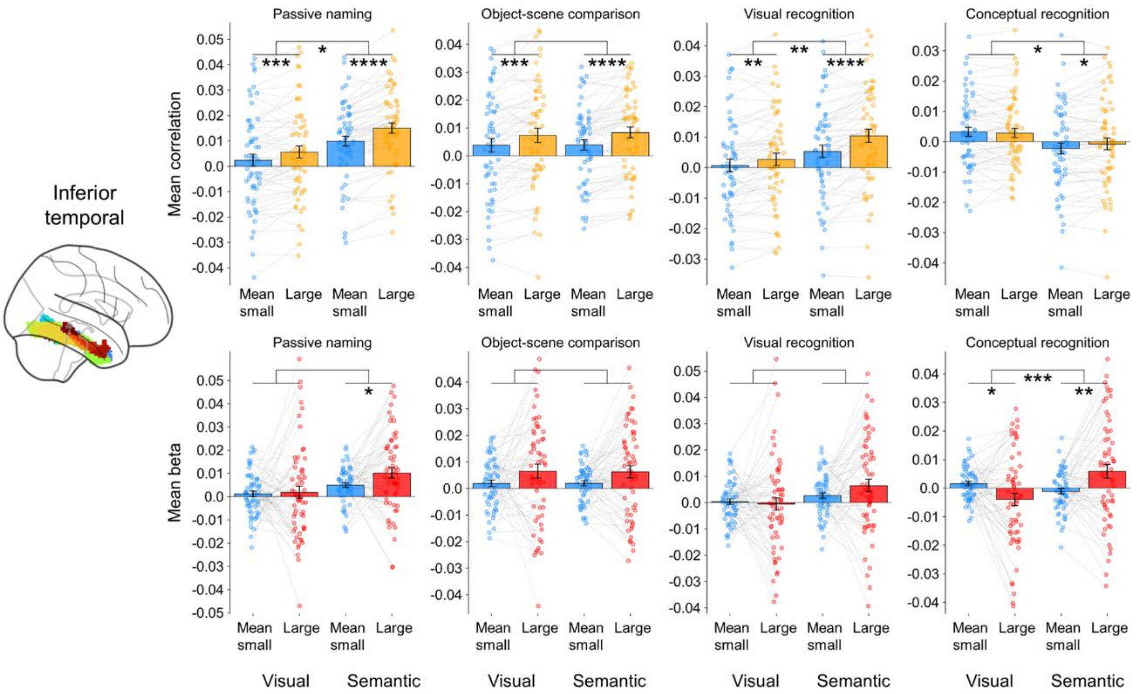


**Figure S3. Multi-scale representational similarity analysis of the four task stages separately.** *The task participants completed included four stages. The main text analyses involved performing representational similarity analyses separately for each of the for stages, and then averaging the resulting correlation coefficients or betas by participant, before submission to paired t-tests and repeated measure analyses of variance (ANOVAs). The data shown here represents the Figure 3 (yellow) and Figure S1 (red) analyses of the inferior temporal lobe data done separately for the (i) passive naming, (ii) object-scene comparison, (iii) visual recognition, and (iv) conceptual recognition tasks; shown in that order from left to right. For details on these tasks' response demands, see below. *, p < .05; **, p < .01; ***, p < .001; ****, p < .0001.*

### 6. Voxel-wise redundancy analysis

Voxel-wise analyses were performed correlating neighboring areas more precisely and to rule out the possibility that the above findings reflect differences in ROIs' shapes or areas' sizes. Gray matter was segmented into 2×2×2 voxel cubes (6×6×6 mm). For each cube, a voxel-wise $NSM_{local}$ was defined and correlated with the $NSM_{local}$ of every other cube within 20 mm. Although not otherwise reported, analyses were also performed with respect to neighbors within 5 mm, 10 mm, or 30 mm, and all the conclusions put forth in the Results remain unchanged. Each cube's correlations with its neighbors were averaged and plotted, which showed patterns consistent with the earlier correlation matrices. Most relevant to the present focus is the ventral red strip in **Figures S4A & S4B**. It corresponds to the inferior temporal gyrus, which had distinctly lower correlations among neighbors than occipital lobe areas. This is consistent with the inferior temporal sub-regions organizing their information processing to operate better as an integrated unit.

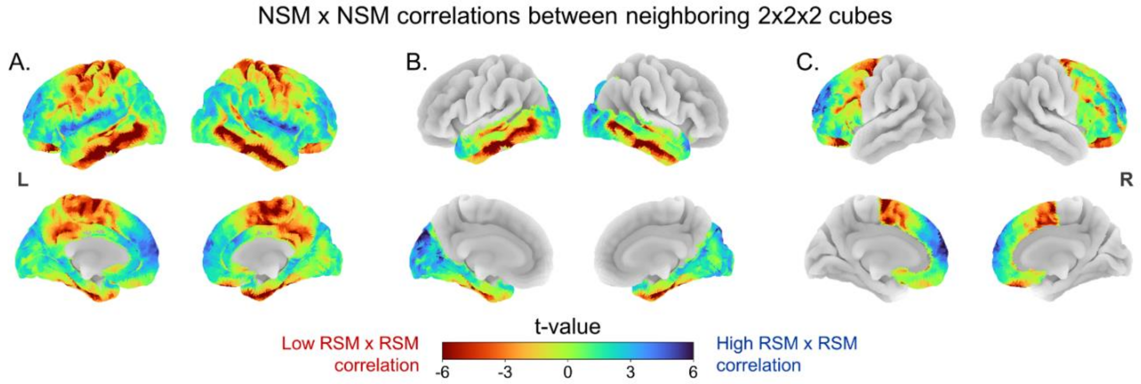


**Figure S4. Cube-wise NSM × NSM correlations.** *Colors are based on the mean NSM × NSM correlation between a 2×2×2 voxel cube at a given location and neighboring cubes within 20 mm. The mean correlations were computed independently for the left and right hemispheres, so no cross-hemisphere interplay is considered, as this could bias medial versus lateral comparisons. For each participant, NSM × NSM correlations were z-standardized across the brain – e.g., if there were just three locations with values 4.0, 3.0, and 2.0, these would be standardized to +1, 0, and -1. The standardized values were averaged across the four stages and submitted to a one-sample t-test, generating the shown t-values. Hence, each location's t-value effectively represents a comparison to other examined locations; this produces slight differences in the t-values across the analyses of (**A.**) the whole neocortex, (**B.**) just the occipital and temporal lobes, and (**C.**) just the prefrontal cortex.*

Interestingly, a pattern emerged in the voxelwise data whereby the orbitofrontal cortex (OFC) displayed lower redundancy than elsewhere in the PFC (**Figure S4C**). Although the main text examination of redundancy focuses on the inferior temporal lobe, this OFC pattern provides an opportunity to further examine the link between redundancy and large-scale coding. Following the presented logic and hypotheses, low redundancy in the OFC suggests the region should also engage in large-scale coding of information. We tested this by returning to the small/large-scale RSA procedures, now comparing the OFC and the remainder of the PFC: These analyses showed that the OFC indeed tends to encode semantic information with large-scale representations, whereas the rest of the PFC does not (**Figure S5**). Although the statistical effect is weaker than the inferior temporal patterns, this is consistent with the logic laid out thus far.

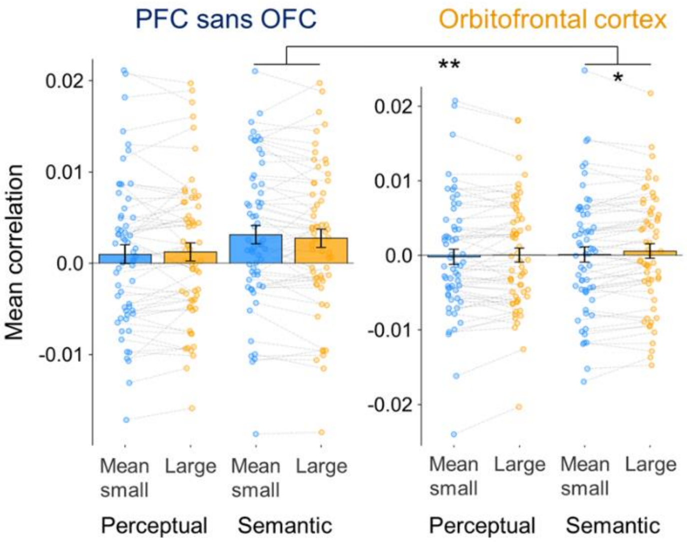


**Figure S5 Examining possible large-scale representation of semantics in the OFC.** *The analysis from main text Figure 3 was conducted for (i) one area defined as the superior frontal gyrus, the middle frontal gyrus, the inferior frontal gyrus, and the anterior cingulate cortex, and (ii) one area defined as just the orbitofrontal gyrus. Paired t-tests were performed for every small/large comparison, and a repeated-measure ANOVA was used for the illustrated across-region interaction effect.*

### 7. Supplemental Methods

#### 7.1. Task design

Participants completed four stages of a study design on object processing, semantics, and memory. In each stage, participants viewed stimuli for the same set of 114 objects, always separated across three runs.

In Stage 1, participants completed trials where they were shown an object (e.g., a tractor) with a label (e.g., "tractor"), and participants used a 4-point scale to rate how accurately the label described the object (1 = "*does not describe the object*"; 4 = "*exact description*"). The task was designed to elicit high ratings (mean rating = 3.60). One week later, participants completed the Stage 2 encoding task (see above) and were asked to return approximately 24hrs (range: 20-28 hours) to complete the Stage 3 and 4, conceptual and perceptual retrieval task, respectively.

For Stage 2, participants were shown a picture of a familiar scene (e.g., a farm) followed by a picture of an object (e.g., a tractor). Participants used a 4-point scale to report how likely it would be to find the object in the scene (1 = "*Very unlikely*", 4 = "*Very likely*"). Each participant completed 114 trials. One-third of trials showed scene-object pairs designed to be incongruent (mean rating = 1.28), one-third showed pairs that were neither congruent nor incongruent (mean rating = 2.11), and one-third showed pairs designed to be congruent (mean rating = 3.67). The scene-object pairs were counterbalanced across participants, so each scene and object appeared in all three conditions, and there were 342 possible scene-object pairs in total.

In the Stage 3 conceptual retrieval task, participants were presented 144 words; 114 were the object labels corresponding to objects shown in the Stage 2 encoding task, and 30 were object labels representing new concepts. For each word, participants indicated whether they had previously seen the object concept during encoding using a 4-point scale (1 = "*definitely new*", 2 = "*probably new*", 3 = "*probably old*", 4 = "*definitely old*"). Participants successfully recognized most objects, and the mean hit rate (3 or 4 response to old images) was 76%. The fMRI analyses would focus only on the 114 old trials, irrespective of participants' responses.

In the Stage 4 visual retrieval task, participants were shown 126 images of objects. Among these, 96 images were of the exact objects shown in the Stage 2 encoding task, 18 images were perceptually similar lures of the original objects (e.g., a blue tractor Stage 2 depicted a red one), and 12 images represented new concepts not seen at the Stage 2 encoding task. Participants responded (i) "old", (ii) "similar", or (iii) "new" to each image, and successfully responded to most images, with a mean old hit rate of 64% and a mean similar hit rate of 49%. The fMRI analyses would focus on the 114 old or similar trials, again irrespective of participants' responses. Preliminary tests showed that the inclusion of miss trials for the conceptual and visual RSA tasks did not influence the patterns of significance in the results.

### 7.2. MRI and preprocessing

#### *7.2.1. MRI acquisition*

MRI data were collected using a General Electric 3T MR750 scanner and an 8-channel head coil. Anatomical images were acquired using a T1-weighted echo-planar sequence (96 slices at

0.9×0.9×1.9 mm$^3$). Functional images were acquired using an echo-planar imaging sequence (repetition time = 2000 ms, echo time = 30 ms, field of view = 19.2 cm, 36 oblique slices with voxel dimensions of 3×3×3 mm). Stimuli were projected onto a mirror at the back of the scanner bore, and responses were recorded using a four-button fiber-optic response box (Current Designs, Philadelphia, PA, USA). Functional resting-state images were collected from the participants using the same parameters (210 volumes, 7 minutes). The BOLD timeseries were resampled into standard space with a spatial resolution of 2×2×2 mm$^3$ or 97×115×97 voxels.

The below descriptions of anatomical and functional preprocessing were automatically generated by fMRIPrep with the express intention that users should copy and paste this text into their manuscripts unchanged.

#### *7.2.2. Anatomical data preprocessing*

A total of two T1-weighted (T1w) images were found within the input BIDS dataset. All of them were corrected for intensity non-uniformity (INU) with N4BiasFieldCorrection [1] distributed with ANTs 2.3.3.[2] The T1w-reference was then skull-stripped with a Nipype implementation of the antsBrainExtraction.sh workflow (from ANTs), using OASIS30ANTs as a target template. Brain tissue segmentation of cerebrospinal fluid (CSF), white-matter (WM), and gray-matter (GM) was performed on the brain-extracted T1w using fast (FSL 6.0.5.1).[3] An anatomical T1w-reference map was computed after registration of two T1w images (after INU-correction) using mri_robust_template (FreeSurfer 7.3.2).[4]

Brain surfaces were reconstructed using recon-all (FreeSurfer 7.3.2),[5] and the brain mask estimated previously was refined with a custom variation of the method to reconcile ANTs-derived and FreeSurfer-derived segmentations of the cortical gray-matter of Mindboggle.[6] Volume-based spatial normalization to one standard space (MNI152NLin2009cAsym) was performed through nonlinear registration with antsRegistration (ANTs 2.3.3), using brain-extracted versions of both T1w reference and the T1w template. The following template was selected for spatial normalization and accessed with TemplateFlow (23.0.0):[7] ICBM 152 Nonlinear Asymmetrical template version 2009c.[8]

#### *7.2.3. Functional data preprocessing*

For each of the seven BOLD runs found per participant (across all tasks and sessions), the following preprocessing was performed. First, a reference volume and its skull-stripped version were generated using a custom methodology of fMRIPrep. Head-motion parameters with respect to the BOLD reference (transformation matrices, and six corresponding rotation and translation parameters) are estimated before any spatiotemporal filtering using MCFLIRT (FSL 6.0.5.1, Jenkinson et al. 2002). BOLD runs were slice-time corrected to 0.972s (0.5 of slice acquisition range 0.00s-1.94s) using 3dTshift from AFNI.[9]

The BOLD time series (including slice-timing correction when applied) were resampled onto their original, native space by applying the transforms to correct for head motion. These resampled BOLD time-series will be referred to as preprocessed BOLD in original space, or just preprocessed BOLD. The BOLD reference was then co-registered to the T1w reference using bbregister (FreeSurfer) which implements boundary-based registration.[10] Co-registration was configured with six degrees of freedom.

Several confounding time series were calculated based on the preprocessed BOLD: framewise displacement (FD), DVARS, and three region-wise global signals. FD was computed using two formulations following Power et al. (absolute sum of relative motions)[11] and Jenkinson (relative root mean squared displacement between affines).[12] FD and DVARS are calculated for each functional run, both using their implementations in Nipype (following the definitions by Power et al.[11]). The three global signals are extracted within the CSF, the WM, and the whole-brain masks. Principal components are estimated after high-pass filtering the preprocessed BOLD time series (using a discrete cosine filter with 128s cut-off) for the two CompCor variants: temporal (tCompCor) and anatomical (aCompCor). tCompCor components are then calculated from the top 2% variable voxels within the brain mask. For aCompCor, three probabilistic masks (CSF, WM, and combined CSF+WM) are generated in anatomical space. The implementation differs from that of Behzadi et al. in that instead of eroding the masks by two pixels on BOLD space, a mask of pixels that likely contain a volume fraction of GM is subtracted from the aCompCor masks. This mask is obtained by dilating a GM mask extracted from the FreeSurfer's aseg segmentation, and it ensures components are not extracted from voxels containing a minimal fraction of GM. Finally, these masks are resampled into BOLD space and binarized by thresholding at 0.99 (as in the original implementation). Components are also calculated separately within the WM and CSF masks. For each CompCor decomposition, the k components with the largest singular values are retained, such that the retained components' time series are sufficient to explain 50% of variance across the nuisance mask (CSF, WM, combined, or temporal). The remaining components are dropped from consideration.

The head-motion estimates calculated in the correction step were also placed within the corresponding confounds file. The confound time series derived from head motion estimates and global signals were expanded with the inclusion of temporal derivatives and quadratic terms for each.[13] Additional nuisance time series are calculated by means of principal components analysis of the signal found within a thin band (crown) of voxels around the edge of the brain, as proposed by Patriat, Reynolds, and Birn.[14] The BOLD time series were resampled into standard space, generating a preprocessed BOLD run in MNI152NLin2009cAsym space. First, a reference volume and its skull-stripped version were generated using a custom methodology of fMRIPrep. All resampling can be performed with a single interpolation step by composing all the pertinent transformations (i.e., head-motion transform matrices, susceptibility distortion correction when available, and co-registrations to anatomical and output spaces). Gridded (volumetric) resampling was performed using antsApplyTransforms (ANTs), configured with Lanczos interpolation to minimize the smoothing effects of other kernels.[15] Non-gridded (surface) resampling was performed using mri_vol2surf (FreeSurfer). Many internal operations of fMRIPrep use Nilearn 0.9.1,[16] mostly within the functional processing workflow. For more details of the pipeline, see the section corresponding to workflows in fMRIPrep's documentation.

#### *7.2.4. Single-trial activity modeling*

Analyses required measuring each voxel's BOLD response in each trial. This was done using first-level general linear models with the Least Squares Separate approach by Mumford et al.,[17] which involves fitting a separate regression for each trial. The regression included a boxcar signal spanning the trial's object presentation period, and this regressor's coefficient represents the trial's BOLD response. The regression also included a boxcar signal covering the presentation time of all other stimuli (i.e., every other object and every scene). Additionally, the linear models included six translation/rotation regressors, three other covariates for head motion (FD, DVARS, and RSMD), and covariates for mean global, white-matter, and cerebrospinal signals. Hence, for all four tasks of each of the 60 participants (240 scans), 114 three-dimensional beta coefficient volumes were defined. These were submitted to RSA.

#### 7.3. Brain area divisions

The analyses also involved organizing a subset of the ROIs into four large anatomical areas based on Brainnetome labels: the *occipital lobe* (22 ROIs; cuneus & occipital gyrus; 139 $cm^3$, including white matter), the *inferior temporal lobe* (32 ROIs; fusiform, parahippocampal, inferior temporal gyri, & anterior temporal lobe [the anterior temporal lobe is not an official Brainnetome label, but is a custom one used by our group]; 138 $cm^3$), the *parietal lobe* (30 ROIs; precuneus, inferior parietal lobule, & super parietal lobule; 192 $cm^3$), and the *prefrontal cortex* (52 ROIs; orbital, inferior, middle, & superior frontal gyri; 288 $cm^3$).

### References

1. Tunison, E., Sylvain, R., Sterr, J., Hiley, V. & Carlson, J. M. No money, no problem: enhanced reward positivity in the absence of monetary reward. *Frontiers in human neuroscience* **13**, 41 (2019).

2. Avants, B. B., Epstein, C. L., Grossman, M. & Gee, J. C. Symmetric diffeomorphic image registration with cross-correlation: evaluating automated labeling of elderly and neurodegenerative brain. *Medical image analysis* **12**, 26–41 (2008).

3. Zhang, Y., Brady, M. & Smith, S. Segmentation of brain MR images through a hidden Markov random field model and the expectation-maximization algorithm. *IEEE transactions on medical imaging* **20**, 45–57 (2001).

4. Reuter, M., Rosas, H. D. & Fischl, B. Highly accurate inverse consistent registration: a robust approach. *Neuroimage* **53**, 1181–1196 (2010).

5. Dale, A. M., Fischl, B. & Sereno, M. I. Cortical surface-based analysis: I. Segmentation and surface reconstruction. *Neuroimage* **9**, 179–194 (1999).

6. Klein, A. *et al.* Mindboggling morphometry of human brains. *PLoS computational biology* **13**, e1005350 (2017).

7. Ciric, R. *et al.* TemplateFlow: FAIR-sharing of multi-scale, multi-species brain models. *Nature Methods* **19**, 1568–1571 (2022).

8. Fonov, V. S., Evans, A. C., McKinstry, R. C., Almli, C. R. & Collins, D. Unbiased nonlinear average age-appropriate brain templates from birth to adulthood. *NeuroImage* **47**, S102 (2009).

9. Cox, R. W. & Hyde, J. S. Software tools for analysis and visualization of fMRI data. *NMR in Biomedicine: An International Journal Devoted to the Development and Application of Magnetic Resonance In Vivo* **10**, 171–178 (1997).

10. Greve, D. N. & Fischl, B. Accurate and robust brain image alignment using boundary-based registration. *Neuroimage* **48**, 63–72 (2009).

11. Power, J. D. *et al.* Methods to detect, characterize, and remove motion artifact in resting state fMRI. *Neuroimage* **84**, 320–341 (2014).

12. Jenkinson, M., Bannister, P., Brady, M. & Smith, S. Improved optimization for the robust and accurate linear registration and motion correction of brain images. *Neuroimage* **17**, 825–841 (2002).

13. Satterthwaite, T. D. *et al.* An improved framework for confound regression and filtering for control of motion artifact in the preprocessing of resting-state functional connectivity data. *Neuroimage* **64**, 240–256 (2013).

14. Patriat, R., Reynolds, R. C. & Birn, R. M. An improved model of motion-related signal changes in fMRI. *Neuroimage* **144**, 74–82 (2017).

15. Lanczos, C. Evaluation of noisy data. *Journal of the Society for Industrial and Applied Mathematics, Series B: Numerical Analysis* **1**, 76–85 (1964).

16. Abraham, A. *et al.* Machine learning for neuroimaging with scikit-learn. *Frontiers in neuroinformatics* **8**, 14 (2014).

17. Mumford, J. A., Turner, B. O., Ashby, F. G. & Poldrack, R. A. Deconvolving BOLD activation in event-related designs for multivoxel pattern classification analyses. *Neuroimage* **59**, 2636–2643 (2012).
